# Performance of deep convolutional neural network approaches and human level in detecting mosquito species

**DOI:** 10.1101/2021.07.23.453554

**Authors:** Rangsan Jomtarak, Veerayuth Kittichai, Theerakamol Pengsakul, Naphop Phatthamolrat, Kaung Myat Naing, Teerawat Tongloy, Santhad Chuwongin, Siridech Boonsang

## Abstract

Recently, mosquito-borne diseases have been a significant problem for public health worldwide. These diseases include dengue, ZIKA and malaria. Reducing disease spread stimulates researchers to develop automatic methods beyond traditional surveillance Well-known Deep Convolutional Neural Network, YOLO v3 algorithm, was applied to classify mosquito vector species and showed a high average accuracy of 97.7 per cent. While one-stage learning methods have provided impressive output in *Aedes albopictus, Anopheles sinensis* and *Culex pipiens*, the use of image annotation functions may help boost model capability in the identification of other low-sensitivity (< 60 per cent) mosquito images for *Cu. tritaeniorhynchus* and low-precision *Ae. vexans* (< 80 per cent). The optimal condition of the data increase (rotation, contrast and blurredness and Gaussian noise) was investigated within the limited amount of biological samples to increase the selected model efficiency. As a result, it produced a higher potential of 96.6 percent for sensitivity, 99.6 percent for specificity, 99.1 percent for accuracy, and 98.1 percent for precision. The ROC Curve Area (AUC) endorsed the ability of the model to differentiate between groups at a value of 0.985. Inter-and intra-rater heterogeneity between ground realities (entomological labeling) with the highest model was studied and compared to research by other independent entomologists. A substantial degree of near-perfect compatibility between the ground truth label and the proposed model (k = 0.950 ± 0.035) was examined in both examinations. In comparison, a high degree of consensus was assessed for entomologists with greater experience than 5-10 years (k = 0.875 ± 0.053 and 0.900 ± 0.048). The proposed YOLO v3 network algorithm has the largest capacity for support-devices used by entomological technicians during local area detection. In the future, introducing the appropriate network model based methods to find qualitative and quantitative information will help to make local workers work quicker. It may also assist in the preparation of strategies to help deter the transmission of arthropod-transmitted diseases.

## Introduction

Mosquito-borne diseases are a significant problem for human health. More than one million cases have been recorded, with an estimated 400,000 deaths in 2018 [1]. These mosquito diseases mostly included dengue fever, zika and malaria, which are prevalent in tropical and subtropical areas. However, they can still spread and cause illness in other areas of the world due to human mobility: globalization, labour movement and transport. In order to minimize morbidity and disease-related mortality, the WHO has urged researchers to further develop more efficient methods that can be used in the entomological field [2].

Aedes genus mosquitoes (*Ae. aegypti, Ae. albopictus*, and *Ae. vaxans*) are pathogens with the primary focus because they can spread human arboviral diseases: dengue, chikungunya, zika and yellow fever [3, 4]. In addition, Anopheles mosquitoes (*An. dirus, An. minimus, An. sinensis* and *An. maculatus*) are vectors for human malaria parasites and can cause a high death rate [5, 6]. Culex mosquitoes can also spread all arboviruses; West Nile virus and human blood parasites (microfilaria and leishmaniae species) [7, 8]. Although the distribution of mosquito vectors is unique to the area, the arthropod-borne pathogens mentioned above are distributed globally by human mobility [9]. Mosquito density and behavior have led us to understand the current effect of disease transmission and spread throughout the region. Procedures for the detection and dissemination of mosquito species require fundamental surveillance. The approach includes highly educated individuals with expertise in the microscopic analysis of insect species [10]. In comparison, it is time-consuming and expensive to complete the monitoring successfully. In addition, the process of preservation is of considerable importance in storing complete and preserved samples of mosquitoes [11, 12]. The deformity of the morphological features of the mosquitoes limits the precision of its classification [13]. Although proposals have been made for the use of high throughput techniques such as PCR, Real-Time PCR and DNA barcode to replace conventional procedures [14-16], the following techniques are not rapid enough and involve qualified molecular-based staff.

Automatic classification tools for mosquito species have been researched to better assist local health personnel. Classification was successfully conducted using an analysis of image characteristics and a flight tone for insects [17-19]. Advancing computerized science offers major classification methods such as artificial intelligence, machine learning, the convolutional neural network (CNN) and the Deep Convolutional Neural Network (DCNN). Image processing is likely to include more than any other method since it can be used to classify insects across their life cycle, including embryos, larvae, pupae and adult stages [20-23]. Previous studies to endorse the hypothesis were to classify and count mosquito eggs in order to better determine the associated population density [24-27]. Any areas of the body, such as wings of insects, have been investigated to classify cryptic mosquito species (as in malaria mosquitoes) [10, 28]. The method, however, faces several obstacles; it is time-consuming and requires human expertise to prepare wing samples. Therefore, the use of the entire body for a feature analysis is assumed to be the most realistic method, because no equipment for sample processing is needed.

Deep learning on the basis of object detection is proportional to classification. In reality, it can also be used to detect both dense freezing and moving objects [29-31]. However, with regard to the concept as mentioned above, the method often overlooks the time-interval lag; between trap environment and microscopic analysis when introduced in actual environments. After making a breakthrough in deep learning; in competition with the ImageNet Wide Scale Visual Recognition Challenge (ILSVRC), target detection has surpassed the state-of-the-art neural network models to reduce the extent of human computational error. Deep learning methods or DCNNs have been developed to classify entomological fields and with high network accuracy. Present high-performance CNN documented its use of Aedes Larvae Recognition, which used a mobile phone-based study of photos captured [26]. However, the model had at least a 30% rate of misclassification. Another application was used to map the distribution of mosquito species using an acoustic recorder on a mobile phone. It is useful for detecting the wing beat of insects [18]. This study helped to quantify the effect of insect density unique to some endemic areas, however the recording method could only be carried out at short distances [18, 19, 32]. Various recording locations of mosquito sound and food-seeking times needed to be validated [17]. Fortunately, high model performance has been proposed, two versions of the neural network You Only Look Once (YOLO v3) have been shown to run faster than other recognition systems because of the network algorithm [31, 33]. The YOLO algorithm is renowned due to its fast object detection speed and high accuracy performance. It solves the issue of object identification in a regression. It then can directly identify the classification probability from the input of images under one CNN and also detect the global data via images from end-to-end training.

This study has been preceded by our previous work that identified gender and also species of field-caught mosquito vectors [34]. The aim of this study is to choose the appropriate model for the classification and position of single mosquito images by comparing two models, respectively YOLO v3 tiny and YOLO v3. The research was undertaken on the basis of the original learning approaches used in the chosen YOLO model. In comparison, small sampling sizes have been used to find acceptable data-augmentation parameters, including rotation, contrast, Gaussian noise and blur. In addition, the degree of acceptance was achieved between the best chosen model and the ground truth, which was also related to independent entomologists. Inter- and intra-human variability has been accomplished by encouraging them to analyze the image sample as being the same as the model test set. Here is to observe whether the model chosen is effective enough to be applied in actual scenarios.

## Materials and Methods

### Mosquito Datasets

In the study, image sets were obtained from two independent open sources (Fig 1). All mosquito images were publicly available from the GitHub repository with the url: https://github.com/jypark1994/MosquitoDL [22]. There are five species of vector mosquitoes including: dengue and zika vectors as *Aedes albopictus* and *Ae. vaxans*; malaria vector as *Anophele*s spp., West Nile virus vector as *Culex pipiens*, Japanese encephalitis vector as *Cu. Tritaeniorhynchus*. As for other species or non-vector mosquitoes, include the: *Ae. dorsalis, Ae. koreikus* and *Cu. inatomii*. The mosquito images were taken from only one source as above to gain the advantage with less variation in image quality. It could benefit the proposed neural network model to reduce the time interval during machine training. In addition, non-mosquito images were obtained from unpublished data: of which included the housefly, stingless bee, and saw-toothed grain beetle.

**Figure 1.**
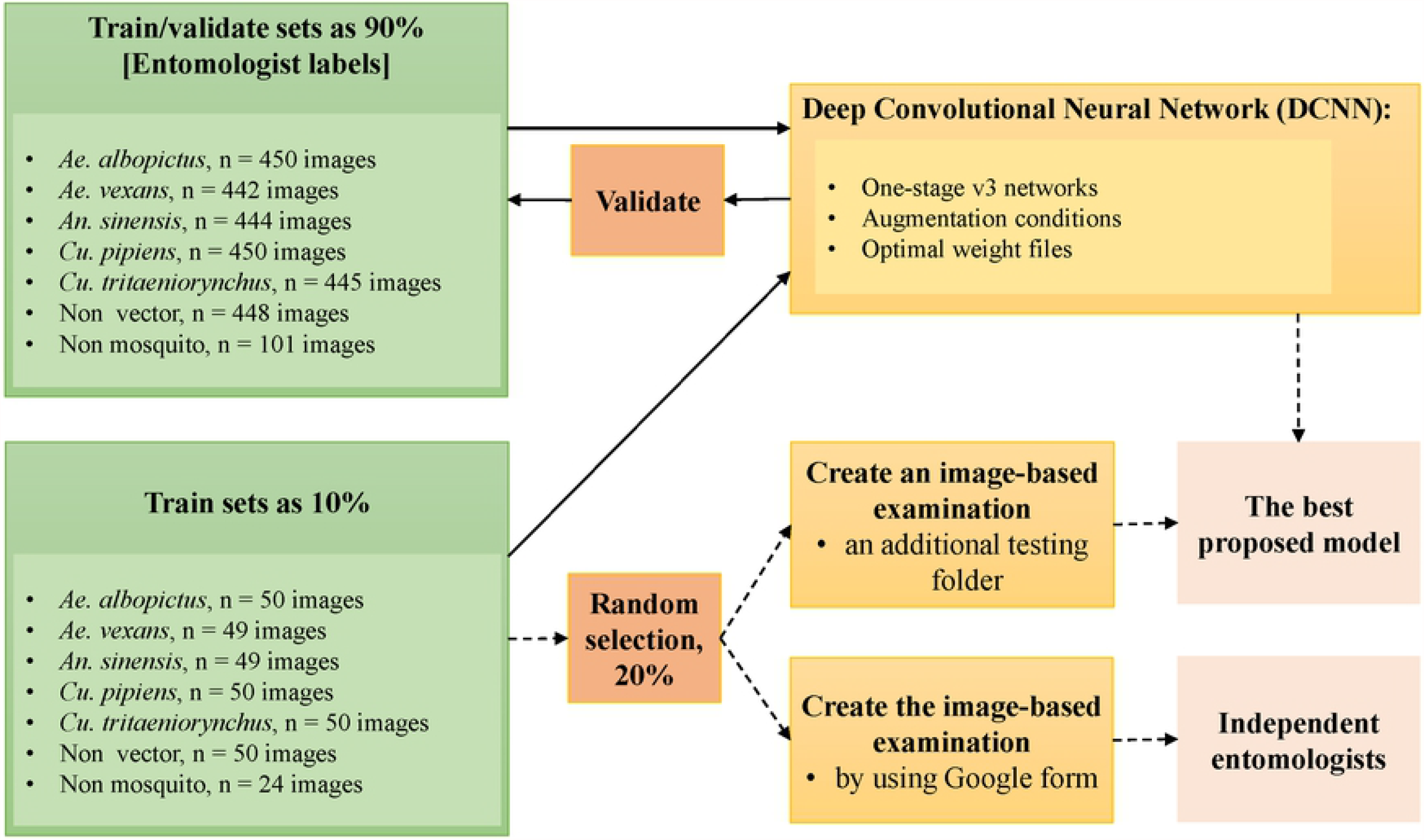
Flowchart of image-set designs used and labeling strategies. 2,780 images obtained from two public-available databases. Main poses were restricted to two-lateral sides but, additional some dorsal- and ventral sides were recruited. Comparison of the model’s performance with the independent entomologists (dash line) happened after training models were achieved (thickness line).

A total of 2,780 images were compiled and divided into training/validation and testing sets. There were five classes of vector species with 2,231 images; and one class for both non-vector and non-mosquito species with 448 and 101 images, respectively. In realistic environments, field-captured mosquitoes had deformations in their body parts and had lost their characteristics. Also, the mosquito variable pose represents the variety of the image. However, the image collection mainly captured side, upper, and ventral views that were mainstreamed for working with machine learning fieldwork [13]. In addition, the pixel densities of each image used in the analysis is compatible with those of the initial resolutions 952 × 1944 and 1920×1080 pixels for the first and second open source. Although the resolution pixels were different in the individual datasets, as seen above, their relative pixel densities were high enough for further training and evaluation of the proposed models. On the basis of data from previous studies, it confirms the concept that the size of image resolution for learning machine learning is at least 320 × 320 pixels [33, 35]. Hence, the use of different image resolutions were used to learn the neural network model.

Of the total image set, the species of mosquitoes and non-mosquitoes is divided into separate directories. These files have been randomly assigned to training/validation and testing sets. They retained a raw test range of 10% for each folder. The remaining images were randomly divided into training/validation or 90% for each folder, the same as before. These insect-specific directories were used to train and validate the proposed model based on a one-stage learning method.

For the learning method inside the proposed model, namely the one-stage learning method, the dataset used under the one-stage method is arranged into a folder. These image sets were responsible for each class of insects and were labelled on the basis of a rectangular box (ground-truth labeling) with a limited potential area per image, a potential region of interest (ROI). The bounding box was used to study whether or not there were any species of mosquitoes. The mark of ground truth was conducted by entomologists under the CiRA CORE program were publicly available from the GitHub repository with the url: https://git.cira-lab.com/cira-medical/cira-mosquitoes-detection [34], based on the species of relative mosquito.

## Development of Deep Neural Networks

The objective of this part was to find the suitable model for classification and localization of every single mosquito within a testing image between Yolo v3 and Yolo tiny-v3 neural network models. These two models studied were implemented to the CiRA CORE program.

In order to guarantee that there was an adequate number and variance of images (because these variables which have an impact on model performance) a distinction between raw images (no annotation of images) and several conditions of the image augmentation was examined (Fig 2). Five conditions were tested, including the default condition at any 4-degree rotational angle increment and 10 percent improvement in brightness/contrast condition, the default and 9-step blur condition, the default and 9-step Gaussian noise condition, and the default and blur and Gaussian noise condition. The conditions-wise comparison was analyzed in relation to the performance of the model. Default condition was obtained by using 45 steps at rotational angles (every 8 degrees) between minimum and maximum [-180 to 180]. Default image contrasts were also modified at every 0.2 stage (with a variance of ± 25 percent) between 0.4 and 1.2. Next, the blur conditions were adjusted for nine steps at each step. The final condition was Gaussian noise, which was corrected for ten steps at each step.

**Figure 2.**
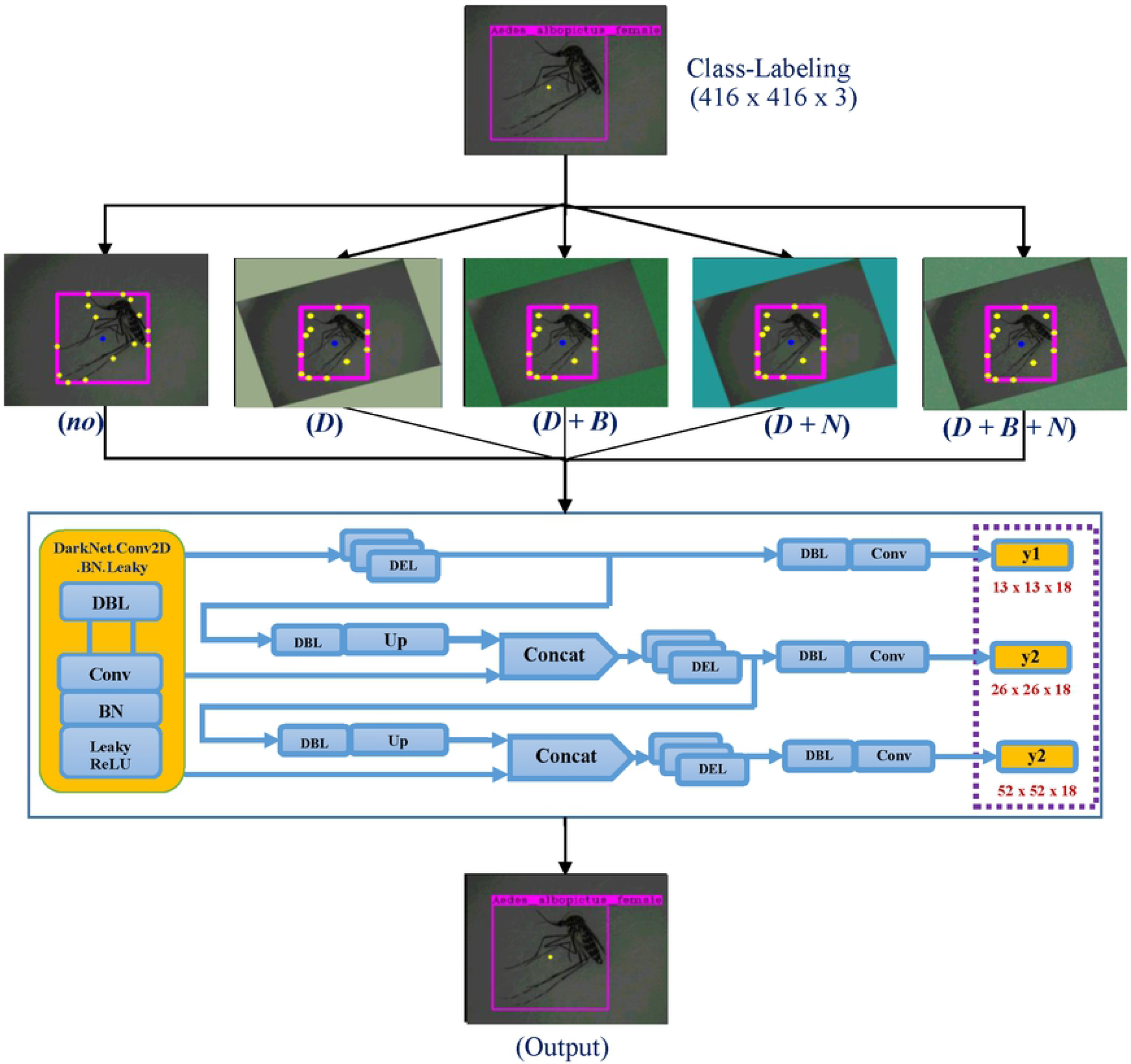
Schematic of labeling and data augmentation conditions with one-stage YOLO v3 architecture model for object detection. For augmentation, “no” is representative for non-augmentation. “***D***” was abbreviated for default (rotational angle and contrasting), “***D + B***” for default and blur, “***D + N***” for default and Gaussian noise, and “***D + B + N***” for default and blur and Gausian noise. A threshold of probability was the confidential value obtained from this equation of *Confidence = Pr(Object) + IOU*^*Truth*^_*Pred*_.

For model training and evaluation, it was run on PyTorch deep learning framework within an Nvidia RTX2070 GPU platform. Learning rates were set at 0.001, which was assumed by the trained weight, reaching optimal accuracy.

To study how effective the chosen model was, it was then compared to human ability. Inter- and intra-human variation in the identification of the tested image sets were assigned.

### Inter and intra-rater variability in human-level

Inter-and intra-human variation was designed to allow observations between model and ground truth registering; relevant to the distribution of the performance of the examiner (entomologist) in the detection of the five species of mosquito vectors. Two independent rounds were completed with a two-month interval. In the first round, 15 independent raters with less than 5 years of experience were anonymously assigned to take the test. 10% % of the test image set was selected and prepared for all raters via Google form (Fig 1). In addition, 25 independent raters were recruited to take part in the second evaluation, which varied from the previous version (images were selected by random). The proposed network model and the performance of the examiners were then observed on the basis of the degree of agreement with the specialist entomologists who labeled the ground-truth as mentioned above.

### Evaluation of model performance

Performance of the proposed models was evaluated by several statistical parameters including: precision, sensitivity, accuracy and specificity [35]. The formulas for these parameters were shown as:

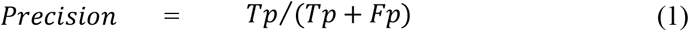

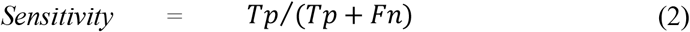

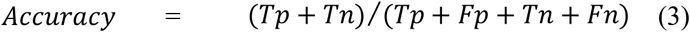

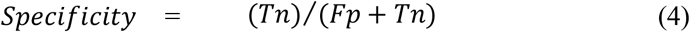

where *Tp* is the number of true positive classifications, *Tn* is the number of true negatives, *Fp* is the number of false positive classifications, *Fn* is the number of false negatives, Actual Positive is the summation between the number of true positive and false negative and Actual Negative is the summation between the number of true negatives and false positives. The precision versus recall curve for every mosquito species was constructed to evaluate the weight performance.

In addition, the performance of the proposed model was assessed by calculating the area under the receiver operating curve (ROC) with 95% confidence intervals (CIs) and the area under the curve (AUC) to determine the accuracy of the models using python. The ROC curve was plotted on the basis of the likelihood value of the 5% increment relative threshold. The 95 percent CIs is measured using a non-parametric bootstrap approach of 1000-fold image re-sampling.

The degree of consensus between the model and the ground truth, as well as the independent entomologists, was studied for exploratory observation by using two operational points, namely sensitivity and specificity. The examiners were not persons who labeled the training images. The test image-set for assessing the performance of both the model and the examiners was the original modal with a 10 % margin parameter in the test static. The degree of consensus in both the proposed model and the examiners was compared to the gold standard; and then calculated by Cohen’s kappa (SPSS software). The Kappa statistics varies from 0 to 1 by following; 0 – 0.20 an agreement equivalent to chance or none (0-4% reliability), 0.21 – 0.39 a minimal agreement (4-15% reliability), 0.40 – 0.59 a weak agreement (15-35 % reliability), 0.60 – 0.79 a moderate agreement (35-63 % reliability), 0.80 – 0.90 a strong agreement (64-81 % reliability), and above 0.90 an almost perfect agreement (82-100 % reliability) [36]. The null hypothesis (*H*_*0*_) was rejected by p-value less than 0.05.

## Results

### Comparison of performance of the network model

The assessment of the two models were investigated based on a threshold probability, *P(t)*, from t_5%_ to t_95%_. The predictions obtained from these models were provided by *P*_*class*_ ≥ *t* [37]. Considering the model’s performance, the capacity of YOLO tiny v3 and YOLO v3 algorithms were compared. The YOLO v3 algorithm showed a higher average accuracy at 97.7% than the YOLO v3 tiny algorithm (Table 1). It has been reported that the rationale of high-performance may be due to the YOLO v3 network having a larger amount of hidden layers and parameters, which is more than any other model version; Tiny-YOLO v3 [31, 33].

**Table 1.** Model-wise comparison. Comparison of the performance of both YOLO v3 versions were studied. Average accuracy by model and also by class was shown.

| Learning<br>method of the<br>network model | Mosquito species |  |  |  |  |  | Average<br>accuracy |
| --- | --- | --- | --- | --- | --- | --- | --- |
|  | *Ae. albopictus* | *Ae. vexans* | *An. sinensis* | *Cu. pipiens* | *Cu. tritaeniorynchus* | Non-vector |  |
| YOLO v3 tiny | 1.000 | 0.946 | 0.970 | 0.980 | 0.923 | 0.997 | **0.969** |
| YOLO v3 | 1.000 | 0.953 | 0.993 | 0.993 | 0.923 | 0.997 | **0.977** |
| **Average**<br>**accuracy** | **1.000** | **0.950** | **0.981** | **0.986** | **0.923** | **0.997** |  |

Class-wise comparison showed a high average detection accuracy of all mosquito species studies, especially in *Ae. albopictus, Cu. pipiens and An. sinensis* as higher than 97per cent. This may be because these three species have unique characteristics, which helped the network model to learn them clearly. Interestingly, the model can be used to detect *Ae. albopictus* at 100% accuracy, which is greater than the detection for any other classes. This is beneficial for exploring the mosquito species in Thailand where dengue is endemic. By contrast, there are similar characters within the images of both *Ae. vexans* and *Cu. tritaeniorhynchus* caught. There was a reason why model performance gave us lower average accuracy than any others. In accordance to the studied models, it may be beneficial in identifying the mosquito vectors for Zika, dengue, malaria and rare pathogens in the tropical region as West Nile virus.

### Selection of data augmentation conditions

The result showed that the YOLO v3 is a state-of-the-art model since it can provide the best high values with parameters including 90.7% of sensitivity, 99.1% of specificity, 97.7% of accuracy and 96.1% of precision (Table 2). Although the one-stage learning methods provide an impressive performance in some mosquito vectors (*Ae. albopictus, An. sinensis* and *Cu. pipiens*), it increases both sample size and their image variation with annotation functions in the fine-tuned network. This could help improve the model capacity in detecting other mosquito images with low sensitivity (< 60per cent) for *Cu. tritaeniorynchus* and low precision for *Ae. vexans* (< 80per cent) (Table 2). The study aimed to increase the model’s performance when compared with non-augmentation conditions. Four data-augmentation conditions were studied including; no augmentation, default, default and blur, and default-blur and noise. It should be noted that the model achieved high performance (accuracy and precision) when augmentation conditions were used. Specifically, the network model with conditions (default + blur, and default + blur + noise) gave us the highest values when compared with other conditions. Finally, the model with optimal augmentation gave us the highest capacity including 96.6 % of sensitivity, 99.6% of specificity, 99.1% of accuracy and 98.1% of precision. Moreover, the AUC under the ROC curve supported the model capacity to distinguish between classes at a value of 0.985 (Fig 3). Hence, annotation functions are very beneficial for training with the proposed model, which has already been fine-tuned. Within the limited amount of biological-samples, it is expected that an enlargement of the sample size and image variation, by using much more annotation strategies, could help solve the problem effectively.

**Table 2.** Comparison of data augmentation conditions for YOLO v3 network model.

| Parameters | Augmentation condition<br>for YOLO v3 model | Mosquito species |  |  |  |  |  | Average |
| --- | --- | --- | --- | --- | --- | --- | --- | --- |
|  |  | *Ae. albopictus* | *Ae. vexans* | *An. sinensis* | *Cu. pipiens* | *Cu. tritaeniorhynchus* | Non-vector |  |
| **Sensitivity** | *No augmentation* | 1.000 | 0.959 | 1.000 | 0.960 | 0.540 | 0.980 | 0.907 |
|  | *default* | 0.940 | 0.837 | 0.939 | 1.000 | 0.680 | 0.920 | 0.886 |
|  | *Default + blur* | 1.000 | 0.959 | 0.898 | 1.000 | 0.700 | 0.980 | 0.923 |
|  | *Default + noise* | 1.000 | 0.959 | 0.959 | 1.000 | 0.860 | 1.000 | 0.963 |
|  | *Default + blur + noise* | 1.000 | 0.939 | 0.980 | 1.000 | 0.920 | 0.960 | 0.966 |
| **Specificity** | *No augmentation* | 1.000 | 0.952 | 0.992 | 1.000 | 1.000 | 1.000 | 0.991 |
|  | *Default* | 1.000 | 0.952 | 0.996 | 0.964 | 1.000 | 1.000 | 0.985 |
|  | *Default + blur* | 1.000 | 0.936 | 1.000 | 1.000 | 1.000 | 0.996 | 0.989 |
|  | *Default + noise* | 0.996 | 0.968 | 1.000 | 1.000 | 1.000 | 1.000 | 0.994 |
|  | *Default + blur + noise* | 1.000 | 0.976 | 1.000 | 1.000 | 1.000 | 1.000 | 0.996 |
| **Accuracy** | *No augmentation* | 1.000 | 0.953 | 0.993 | 0.993 | 0.923 | 0.997 | 0.977 |
|  | *Default* | 0.990 | 0.933 | 0.987 | 0.970 | 0.946 | 0.987 | 0.969 |
|  | *Default + blur* | 1.000 | 0.940 | 0.983 | 1.000 | 0.950 | 0.993 | 0.978 |
|  | *Default + noise* | 0.997 | 0.966 | 0.993 | 1.000 | 0.977 | 1.000 | 0.989 |
|  | *Default + blur + noise* | 1.000 | 0.970 | 0.997 | 1.000 | 0.987 | 0.993 | 0.991 |
| **Precision** | *No augmentation* | 1.000 | 0.797 | 0.961 | 1.000 | 1.000 | 1.000 | 0.960 |
|  | *Default* | 1.000 | 0.774 | 0.979 | 0.847 | 1.000 | 1.000 | 0.933 |
|  | *Default + blur* | 1.000 | 0.746 | 1.000 | 1.000 | 1.000 | 0.980 | 0.954 |
|  | *Default + noise* | 0.980 | 0.855 | 1.000 | 1.000 | 1.000 | 1.000 | 0.972 |
|  | *Default + blur + noise* | 1.000 | 0.885 | 1.000 | 1.000 | 1.000 | 1.000 | 0.981 |

**Figure 3.**
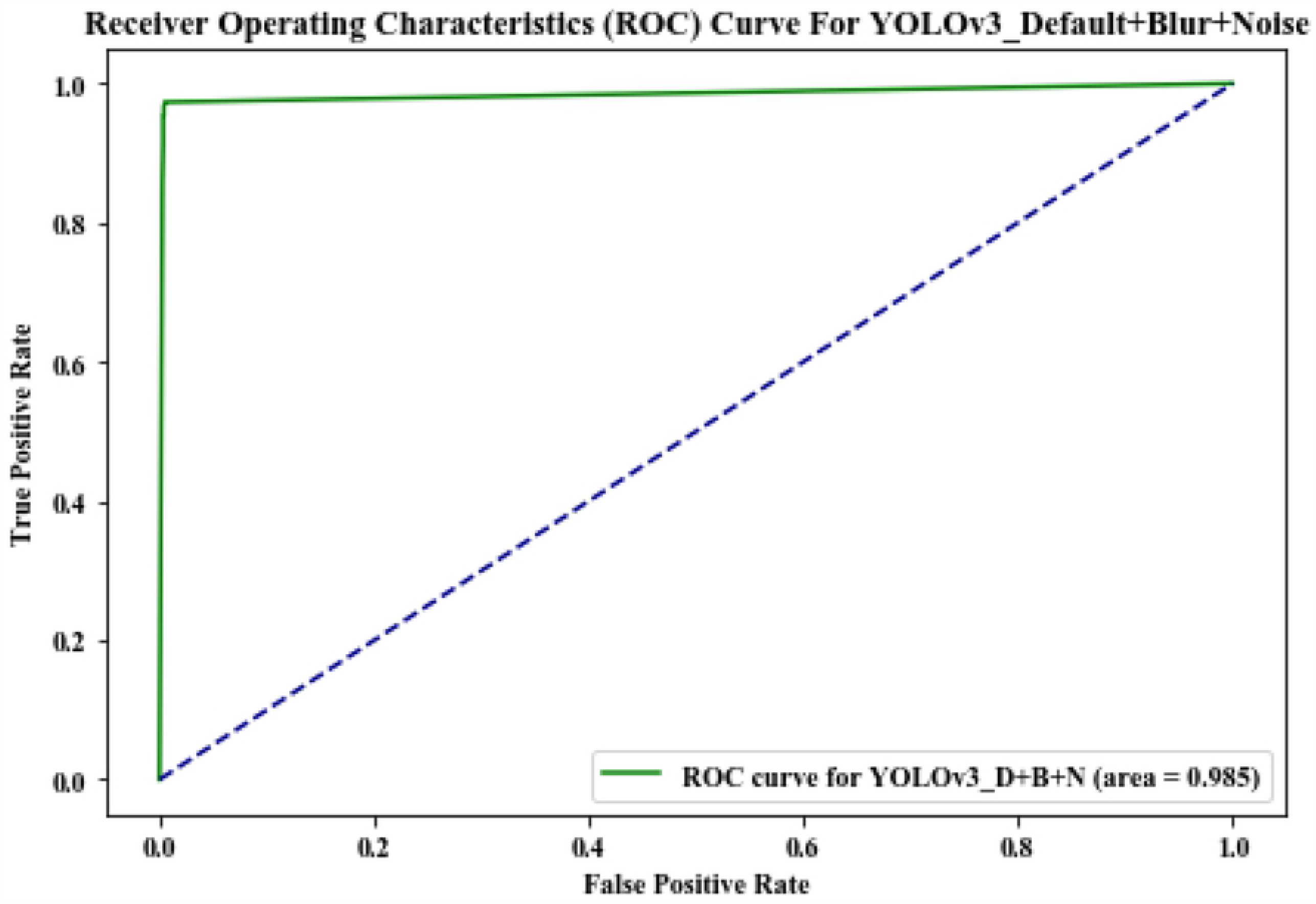
ROC curve of the YOLO v3 model with optimal augmentation condition.

### Inter- and intra-rater variability

Evaluation of performance was studied at the level of consensus of the ground truth (entomological labeling); with the best model and connected to other independent entomologists. Here, determine whether the model chosen was successful enough to be applied in a specific environment. 10 % of the test datasets were randomly chosen and used for blind testing by all human investigators; as inter-and intra-human variants. Approximately one month after the first evaluation, 10 percent of the image re-selection from the test collection was circulated through Google form to the same examiners for more intra-and inter-examination variability.

In the first test, the significant agreement level between the ground truth labeling and the proposed model (*k = 0*.*950 ±* 0.035) and with entomologists, who had more than 5-10 years if experience was perfect degrees (*k = 0*.*875 ± 0*.*053* and *0*.*900 ± 0*.*048*), as measured by Cohen’s kappa (Fig 4a). The results reflect the similar visualities between the DCNN model studied and in humans. Significantly, there was clear evidence in the agreement values for the independent entomologists, with less than five-years of experience, showed a lower degree when compared to the experienced entomologists. The second test, the agreement level values tended to be similar to the first test, except for in subjects s2_3 who got higher scores than the first test from intermediate (*k = 0*.*700 ± 0*.*041*) to perfect levels (*k = 0*.*875 ± 0*.*053*) (Fig 4b). Inter- and intra-human variability confirmed the requirement for long-term training in younger individuals to be specialists. Automatic devices, on the other hand, can achieve excellent performance based on the size of the sample being inputted. The latter emphasizes automated devices that can be used to assist human capabilities and are to be regarded. As there is a reduction in the interest in entomology and the production of human expertise requires a great deal of time, automated systems, on the other hand, have shown that they perform significantly well. As a result, people have become more interested in automatic systems, although the ability of the system does not rely on its processing time, but rather depends on data to learn the model within a time-limited frame.

**Figure 4.**
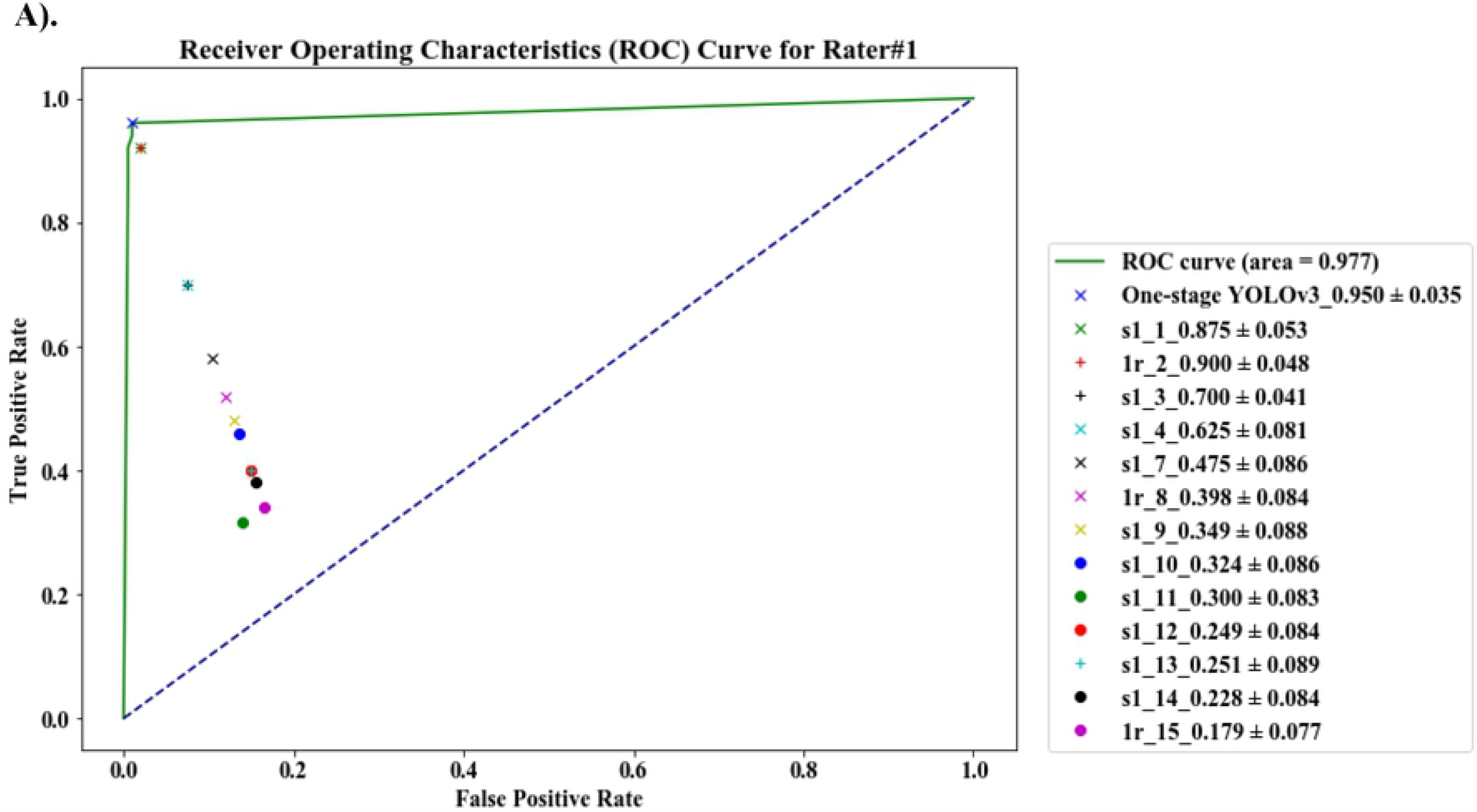
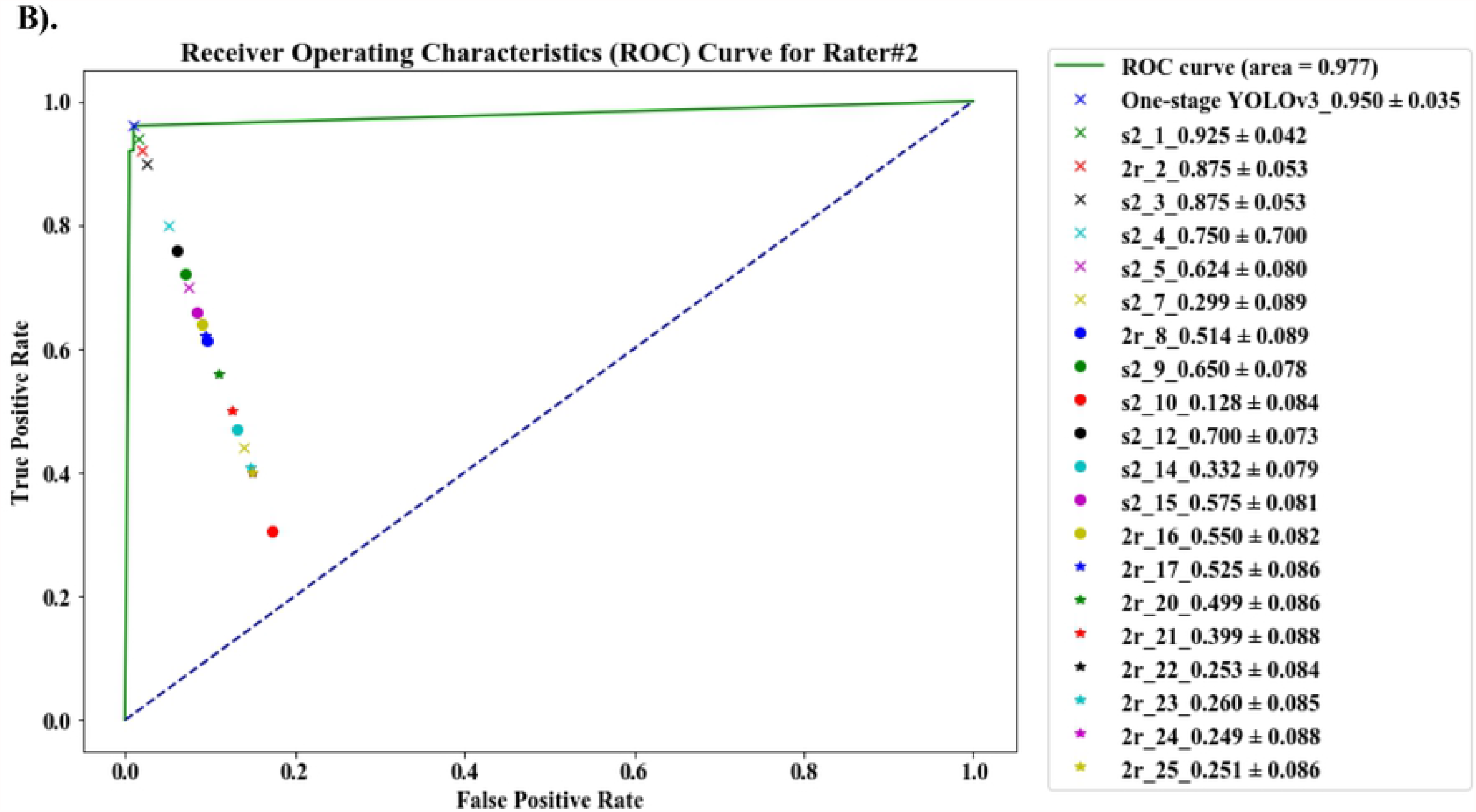
ROC curve with agreement level of the proposed model and the independent-entomologists. ***A*** represents the first test and ***B*** the second test. **S1_1** represents subject-1, who joined the first test and **S2_1** for subject-1 who joined both tests. **1r_2** represents subject-2, who joined the first test only and **2r_2** represents subject-2, who joined only the second test.

In addition, the higher AUC value (0.977) under the ROC curve seen statically indicates some values of average sensitivity and average specificity between the first and second human variability studies, especially in entomologists with less than 5 years of experience. With a short period of time for training between humans and DCNN, the network model can be expected to be useful for further application in remote areas where there is a shortage of expert entomologists.

## Discussion

The DCNN suggested in the research study resulted in high success in the identification of mosquito species that can spread several arthropod-borne pathogens, including dengue virus, ZIKA virus, West Nile virus and malaria parasite. The most prevalent species of mosquito with the special characteristics used enables the system to operate with greater precision, specificity and sensitivity than 96 per cent, resulting in higher accuracy relative to other mosquitoes [20, 22, 38, 39]. Previous work suggested that a larger data set could boost model efficiency for uncommon image classes [37]. In this study, when data augmentations were done; such as combination of rotation, contrasting, Gaussian noise and blur conditions, their model’s performance, it showed outstanding value to non-augmentation which is consistent with the previous report [22]. Furthermore, image sets with different resolution pixels and their illumination may sometimes improve the overall detection capability of the model [37]. The research model demonstrated good success in distinguishing between mosquito and non-mosquito, mosquito vector and non-vector with achieved high accuracy and ROC AUC of approximately 0.985 in the model-wise relation (Fig 3). Consider it scalable and fast with the YOLO v3 algorithm compared to other detection methods such as DPM, R-CNN, and Darknet-19 [31, 33]. The proposed model allowed technicians to detect the object very quickly. This model can be useful in the detection of species of mosquitoes in remote areas which are likely to have a large number of mosquitoes collected from mosquito traps. Object identification predictions can also be extended to entomologically related work, as all organisms could be identified with high confidence using the proposed network model.

Levels of disagreements between the proposed network obtained and the ground truth labeling are connected to inter- and intra-rater variability in the entomological examination. The variability examination in humans was studied by using a two-independent imageset. Two-animal species (*Ae. vexans* and *Cu. tritaeniorynchus*), of particularly high variability. Although these two animals are different in genus-level and differ in their natural characteristics; the difficulty, in both the proposed model and inter- and intra-raters, to distinguish them may be relevant to the image’s quality happening during data collection. For example, the difference in focus quality may make it difficult to label datasets and train models [40]. Although computational modeling has had a significant influence on both clinical and science work, more enhancements are needed. This requires (a) a large number of training details, and (b) a new methodological architecture to be learned to manage images collected from various scanners or cameras [41]. Using the same basic type of camera property and stereo microscope to capture the mosquito image could help promote further deployment of the embedded device network concept in remote areas elsewhere, without re-training the data prior to use in real-time scenarios.

## Conclusion

The proposed YOLO v3 network algorithm provides great potential for rapid screening and support devices for entomological technicians, especially during mosquito identification. According to inter-and intra-human variability, the experiment has been accomplished by encouraging them to analyze the image sample as being the same as the model test set. In the future, qualitative and quantitative devices based on the best network model will help to make it easier for local workers to perform quicker and to prepare strategies related to the advanced prevention of mosquito-borne diseases.

## Acknowledgements

We are grateful to Research Seed Grant for New Lecturers [KREF186110] and Thailand Research Fund [RE-KRIS/016/64], who have provided financial support for the research project. We also thank the College of Advanced Manufacturing Innovation, King Mongkut’s Institute of Technology, Ladkrabang who have provided the deep learning platform and software to support the research project.

## Data availability

The data that support the findings of this study are available upon requested to the corresponding author.

## Author contributions

**Conceptualization:** Rangsan Jomtarak, Siridech Boonsaeng & Santhad Chuwongin

**Data collection:** Natthaphop Phatthamolrat, Theerakamol Pengsakul & Kaung Myat Naing

**Funding acquisition:** Veerayuth Kittichai

**Investigation:** Veerayuth Kittichai & Theerakamol Pengsakul

**Methodology:** Veerayuth Kittichai

**Project administration:** Natthaphop Phatthamolrat

**Software:** Teerawat Tongloy, Santhad Chuwongin, Siridech Boonsaeng

**Supervision:** Rangsan Jomtarak

**Writing – original draft:** Rangsan Jomtarak and Veerayuth Kittichai

**Writing – review & editing:** Veerayuth Kittichai, Theerakamol Pengsakul, Santhad Chuwongin, & Siridech Boonsang

## References

1. WHO. World malaria report. WHO. 2019.

2. WHO. Global Vector Control Response 2017-2030—Background Document to Inform Deliberations during the 70th Session of the World Health Assembly. WHO. 2017:47.

3. Monteiro FJC, Mourao FRP, Ribeiro ESD, Rego M, Frances P, Souto RNP, et al. Prevalence of dengue, Zika and chikungunya viruses in Aedes (Stegomyia) aegypti (Diptera: Culicidae) in a medium-sized city, Amazon, Brazil. Rev Inst Med Trop Sao Paulo. 2020;62:e10.

4. Paixao ES, Teixeira MG, Rodrigues LC. Zika, chikungunya and dengue: the causes and threats of new and re-emerging arboviral diseases. BMJ Glob Health. 2018;3(Suppl 1):e000530.

5. Ouedraogo AL, Bousema T, Schneider P, de Vlas SJ, Ilboudo-Sanogo E, Cuzin-Ouattara N, et al. Substantial contribution of submicroscopical Plasmodium falciparum gametocyte carriage to the infectious reservoir in an area of seasonal transmission. PLoS One. 2009;4(12):e8410.

6. Bousema T, Drakeley C. Epidemiology and infectivity of Plasmodium falciparum and Plasmodium vivax gametocytes in relation to malaria control and elimination. Clin Microbiol Rev. 2011;24(2):377–410.

7. Colpitts TM, Conway MJ, Montgomery RR, Fikrig E. West Nile Virus: biology, transmission, and human infection. Clin Microbiol Rev. 2012;25(4):635–48.

8. Aboagye-Antwi F, Kwansa-Bentum B, Dadzie SK, Ahorlu CK, Appawu MA, Gyapong J, et al. Transmission indices and microfilariae prevalence in human population prior to mass drug administration with ivermectin and albendazole in the Gomoa District of Ghana. Parasit Vectors. 2015;8:562.

9. Sriwichai P, Karl S, Samung Y, Kiattibutr K, Sirichaisinthop J, Mueller I, et al. Imported Plasmodium falciparum and locally transmitted Plasmodium vivax: cross-border malaria transmission scenario in northwestern Thailand. Malar J. 2017;16(1):258.

10. Yang HP, Ma CS, Wen H, Zhan QB, Wang XL. A tool for developing an automatic insect identification system based on wing outlines. Sci Rep. 2015;5:12786.

11. Taai K, Harbach RE, Aupalee K, Srisuka W, Yasanga T, Otsuka Y, et al. An effective method for the identification and separation of Anopheles minimus, the primary malaria vector in Thailand, and its sister species Anopheles harrisoni, with a comparison of their mating behaviors. Parasit Vectors. 2017;10(1):97.

12. Dowell FE, Noutcha AE, Michel K. Short report: The effect of preservation methods on predicting mosquito age by near infrared spectroscopy. Am J Trop Med Hyg. 2011;85(6):1093–6.

13. Daniel da Silva Motta RB, Alex Santos and Frank Kirchner. Use of Artificial Intelligence on the Control of Vector-Borne Diseases, Vectors and Vector-Borne Zoonotic Diseases: IntechOpen; 2018. Available from: https://www.intechopen.com/books/vectors-and-vector-borne-zoonotic-diseases/use-of-artificial-intelligence-on-the-control-of-vector-borne-diseases https://api.intechopen.com/chapter/pdf-download/64098.pdf.

14. Rochlin I, Santoriello MP, Mayer RT, Campbell SR. Improved high-throughput method for molecular identification of Culex mosquitoes. J Am Mosq Control Assoc. 2007;23(4):488–91.

15. Kothera L, Byrd B, Savage HM. Duplex Real-Time PCR Assay Distinguishes Aedes aegypti From Ae. albopictus (Diptera: Culicidae) Using DNA From Sonicated First-Instar Larvae. J Med Entomol. 2017;54(6):1567–72.

16. Shahhosseini N, Kayedi MH, Sedaghat MM, Racine T, G PK, Moosa-Kazemi SH. DNA barcodes corroborating identification of mosquito species and multiplex real-time PCR differentiating Culex pipiens complex and Culex torrentium in Iran. PLoS One. 2018;13(11):e0207308.

17. Arthur BJ, Emr KS, Wyttenbach RA, Hoy RR. Mosquito (Aedes aegypti) flight tones: frequency, harmonicity, spherical spreading, and phase relationships. J Acoust Soc Am. 2014;135(2):933–41.

18. Haripriya Mukundarajan FJHH, Erica Araceli Castillo, Cooper Newby, Manu Prakash. Using mobile phones as acoustic sensors for high-throughput mosquito surveillance. eLife. 2017:1–26.

19. Menda G, Nitzany EI, Shamble PS, Wells A, Harrington LC, Miles RN, et al. The Long and Short of Hearing in the Mosquito Aedes aegypti. Curr Biol. 2019;29(4):709–14 e4.

20. Motta D, Santos AAB, Winkler I, Machado BAS, Pereira D, Cavalcanti AM, et al. Application of convolutional neural networks for classification of adult mosquitoes in the field. PLoS One. 2019;14(1):e0210829.

21. Adam Goodwin MG, Tristan FORD, Laura SCAVO, Jewell BREY, Collyn HEIER, Nicholas J. DURR, AND SOUMYADIPTA ACHARYA. Development of a low-cost imaging system for remote mosquito surveillance. Biomedical Optics Express. 2020;11(5).

22. Park J, Kim DI, Choi B, Kang W, Kwon HW. Classification and Morphological Analysis of Vector Mosquitoes using Deep Convolutional Neural Networks. Sci Rep. 2020;10(1):1012.

23. Kim K, Hyun J, Kim H, Lim H, Myung H. A Deep Learning-Based Automatic Mosquito Sensing and Control System for Urban Mosquito Habitats. Sensors (Basel). 2019;19(12).

24. Shumkov MA. [Methods of detection of Aedes mosquito eggs in the soil]. Med Parazitol (Mosk). 1966;35(5):615–7.

25. Siti Azirah Asmai MNDMZ, Abdul Syukor Mohamad Jaya, Ahmad Fadzli Nizam Abdul Rahman, Zuraida Binti Abal Abas Mosquito Larvae Detection using Deep Learning. International Journal of Innovative Technology and Exploring Engineering (IJITEE). 2019;8(12):804–9.

26. Alejandra Sanchez Ortiz MNM, Henrik Tünnermann, Toya Teramoto, Hayaru Shouno. Mosquito Larva Classification based on a Convolution Neural Network. Int’l Conf Par and Dist Proc Tech and Appl 2018. p. 320–5.

27. Antonio Arista-Jalife MN, Zaira Garcia-Nonoal, Daniel Robles-Camarillo, Hector Perez-Meana, Heriberto Antonio Arista-Viveros. Aedes mosquito detection in its larval stage using deep neural networks. Knowledge-Based Systems,. 2020;189.

28. Lorenz C, Ferraudo AS, Suesdek L. Artificial Neural Network applied as a methodology of mosquito species identification. Acta Trop. 2015;152:165–9.

29. Zhang X, Yang W, Tang X, Liu J. A Fast Learning Method for Accurate and Robust Lane Detection Using Two-Stage Feature Extraction with YOLO v3. Sensors (Basel). 2018;18(12).

30. Unver HM, Ayan E. Skin Lesion Segmentation in Dermoscopic Images with Combination of YOLO and GrabCut Algorithm. Diagnostics (Basel). 2019;9(3).

31. Joseph Redmon SD, Ross Girshick, Ali Farhadi. You Only Look Once: Unified, Real-Time Object Detection. arXiv:150602640 [csCV]. 2016.

32. Albert JT, Kozlov AS. Comparative Aspects of Hearing in Vertebrates and Insects with Antennal Ears. Curr Biol. 2016;26(20):R1050–R61.

33. Joseph Redmon AF. YOLOv3: An Incremental Improvement. arXiv:180402767 [csCV]. 2018.

34. Kittichai V, Pengsakul T, Chumchuen K, Samung Y, Sriwichai P, Phatthamolrat N, et al. Deep learning approaches for challenging species and gender identification of mosquito vectors. Sci Rep. 2021;11(1):4838.

35. Wang Q, Bi S, Sun M, Wang Y, Wang D, Yang S. Deep learning approach to peripheral leukocyte recognition. PLoS One. 2019;14(6):e0218808.

36. McHugh ML. Interrater reliability: the kappa statistic. Biochem Med (Zagreb). 2012;22(3):276–82.

37. Christian Matek SS, Karsten Spiekermann, and Carsten Marr. Human-level recognition of blast cells in acute myeloid leukaemia with convolutional neural networks. Nature Machine Intelligence volume. 2019;1:538–44.

38. Medronho RA, Camara VM, Macrini L. Classification of containers with Aedes aegypti pupae using a Neural Networks model. PLoS Negl Trop Dis. 2018;12(7):e0006592.

39. Kazushige Okayasu KY, Masataka Fuchida and Akio Nakamura. Vision-Based Classification of Mosquito Species: Comparison of Conventional and Deep Learning Methods. Applied sciences. 2019;9(3935).

40. Kohlberger T, Liu Y, Moran M, Chen PC, Brown T, Hipp JD, et al. Whole-Slide Image Focus Quality: Automatic Assessment and Impact on AI Cancer Detection. J Pathol Inform. 2019;10:39.

41. Chan HP, Samala RK, Hadjiiski LM, Zhou C. Deep Learning in Medical Image Analysis. Adv Exp Med Biol. 2020;1213:3–21.

